# Nanogap Solid-State Single-Molecule Detection at Mars, Europa, and Microgravity Conditions

**DOI:** 10.1101/2024.02.29.582359

**Authors:** José L. Ramírez-Colón, Emma Johnson, Daniel Duzdevich, Sam Lee, Jason Soderblom, Maria T. Zuber, Masateru Taniguchi, Takahito Ohshiro, Yuki Komoto, Christopher E. Carr

## Abstract

Solid-state nanogap systems are an emerging technology for *in-situ* life detection due to their single-molecule resolution of a wide range of biomolecules, including amino acids and informational polymers, at the parts per billion to trillion level. By targeting the abundance distributions of organic molecules, this technology is a candidate for detecting ancient and extant life and discriminating between biotic and abiotic organics on future planetary missions to Mars and icy moons such as Enceladus and Europa. A benchtop system developed at Osaka University has a proven ability to detect and discriminate among single amino acids, RNA, and DNA using nanogap chips. The Electronic Life-detection Instrument for Enceladus/Europa (ELIE) prototype was subsequently developed to make this technology viable for space instrumentation through the simplification of electronics, reduction of size and weight, and automation of gap formation. Initial ground testing using a manually formed nanogap with the first ELIE prototype detected the amino acid L-proline. However, this manual adjustment approach posed limitations in maintaining a consistent gap size. To address this challenge, we integrated an automated piezo actuator to enable real-time gap control, permitting single-molecule identification of a target amino acid, L-proline, under reduced gravity (*g*), including Mars (*g* = 0.378), Europa or Lunar (*g* = 0.166), and microgravity conditions (*g* = 0.03-0.06), as validated through parabolic flight testing. Power supply noise and experimental constraints of the experiment design limited data collection to short segments of good-quality data. Nevertheless, the subsequent analysis of detected events within these segments revealed a consistent system performance and a controlled gap size across the different accelerations. This finding highlights the system’s resilience to physical vibrations. Future goals are to progress the instrument towards technology readiness level 4 with further reductions of size and mass, lower noise, and additional system automation. With further development, ELIE has the potential to be an autonomous and sensitive single-molecule detection instrument for deployment throughout the solar system.

## 1. Introduction

Technological advancements have brought the exploration of Ocean Worlds, like Enceladus and Europa, within reach, positioning these potentially habitable moons as primary candidates in the quest for life beyond Earth [1–5]. One approach to searching for life on these moons could involve seeking signatures of life at the molecular level. Life as we know it relies on specific organic molecules, including a group of 20 amino acids that constitute the building blocks of proteins [6]. More than half of these amino acids have been found in meteorites, suggesting they can be formed abiotically and do not necessarily originate on Earth [7–9]. However, the distribution of amino acids detected in meteorites results in an excess of the simplest amino acid (glycine) and in mostly racemic mixtures of the amino acids present [10, 11]. This differs from systems modified by biotic processes where more complex amino acids and homochirality (e.g., preference for L-amino acids) are observed [12, 13]. This distribution of amino acids, both in type and chirality, could serve as a biosignature to distinguish between a biological and non-biological origin of the molecules.

Seeking these signatures of life at a molecular level on Ocean Worlds requires ultra-sensitive instrumentation, capable of discerning among amino acids expected at low abundances [14]. Such an instrument must also be adaptable to different types of biochemistries, meet the mass, power, and volume constraints required for space missions, and withstand harsh space conditions like launch vibrations, radiation, temperature, pressure, and microgravity. Solid-state nanogaps have emerged as attractive single-molecule detection platforms for biomolecule analysis, including nucleic acids and proteins [15–17], nucleobases and amino acids [18], and other macro-molecules [19, 20]. Solid-state nanopores and nanogaps are nanoscale structures fabricated within solid-state materials (e.g., silicon nitride, graphene), with nanopores being tiny holes ranging from a few to hundred nanometers in size and nanogaps consisting of closely spaced electrodes with nanometer-sized gaps between them. Their sub-nanometer precision and pore size rangeability make them ideal sensors for low-concentration sample detection and highly sensitive to molecules of different sizes on the nanometer scale.

One such instrument, the Electronic Life Detection Instrument for Enceladus/Europa (ELIE), utilizes nanogap sensors that work based on quantum electronic tunneling to detect single molecules (Figure 1). These sensors have previously been demonstrated to detect and discriminate among single amino acids [21], nucleobases [22], and nucleic acids [23]. ELIE is designed to primarily target molecular biosignatures (e.g., amino acids and informational polymers) in future *insitu* life detection missions. The first low-technology readiness level (TRL) ELIE prototype proved the detection of the amino acid L-proline and demonstrated an extrapolated sensitivity of 1 nM [24]. However, this prototype required manual adjustments to maintain the nanogap, which could only be sustained for short intervals.

**Figure 1.**
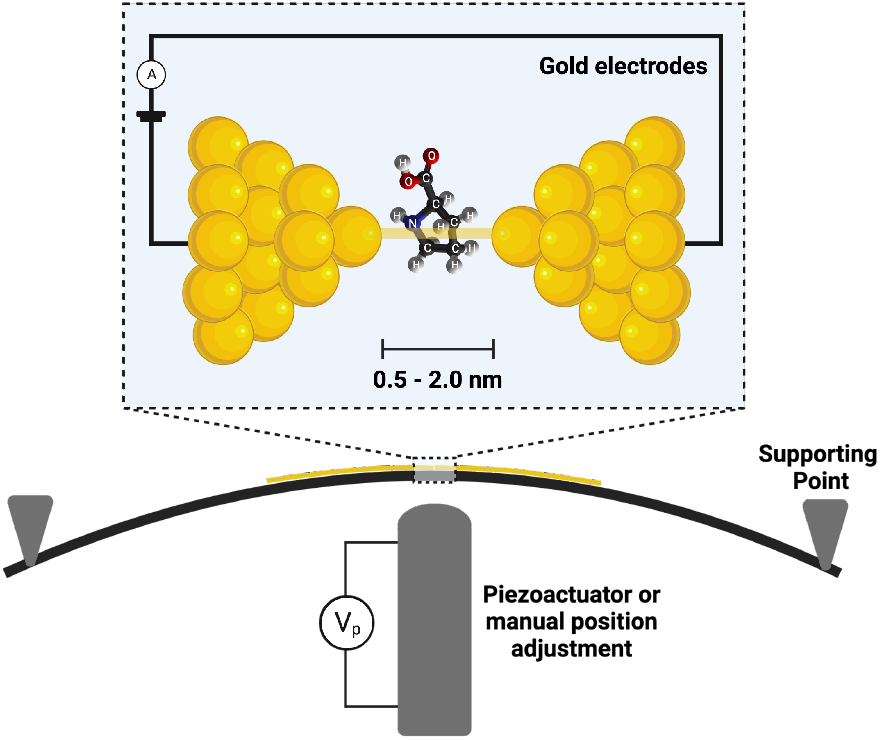
Single-molecule detection with a quantum electronic tunneling nanogap sensor, using the Mechanically Controlled Break Junction method.

Considering the limitation of this first prototype, we have developed a next-generation prototype, which integrates an automated piezo actuator for gap control. Furthermore, we tested the prototype during a parabolic flight to quantify the impact of altered *g* level and vibration on the single-molecule detection of the amino acid, L-proline. In parallel, we performed control experiments on the ground to assess potential differences in the performance of ELIE related to the instrument’s noise, event detection, and event characterization. We demonstrate that a solid-state single-molecule device works in microgravity and continues to function after multiple changes in gravitational force. These findings provide evidence of the feasibility of integrating ELIE into future *in-situ* life detection missions, advancing our pursuit of potential extraterrestrial life within our solar system.

## 2. Methods

### ELIE Instrument Hardware

The first-generation ELIE prototype originated from a bench-top nanogap system (herein referred to as AXN) built by the Taniguchi laboratory at Osaka University, as reported in Tsutsui *et al*. 2008, Ohshiro *et al*. 2012, Ohshiro *et al*. 2014, and Ohshiro *et al*. 2018 [21–23, 25]. This system comprises a picoammeter, a National Instruments computer, a custom Faraday cage enclosing stepping piezo motors, and a jig structure holding a nanogap chip, a piezo controller, and battery banks. With the goal of reducing the mass and volume of this benchtop system, the first-generation ELIE prototype was built, consisting of a low-noise, high-bandwidth amplifier (Chimera Instruments, VC100, 8 pA RMS at 100 kHz, with sampling up to 4 MHz), a laptop for data collection, and a Faraday cage holding an amplifier head stage, a nanogap chip, and a manual micrometer. The manual adjustment approach posed limitations in maintaining a consistent gap size. In this next-generation prototype, an automated piezo actuator (PI N-381) with a piezo controller (PI E-861) was integrated into the design to enable real-time gap control (Figures 2a and 2b). The low-noise amplifier and the Faraday cage were mounted onto a 24” x 24” baseplate using 3M™ Dual Lock™ (SJ3560). The nanogap chip (Figure 2c) consists of a silicon substrate coated with a thin polyimide layer and patterned gold nanojunctions on top. A detailed description of the fabrication process is described in Tsutsui *et al*. 2008 [25]. A polydimethylsiloxane cover was integrated on top of the chip for sample containment and retention.

**Figure 2.**
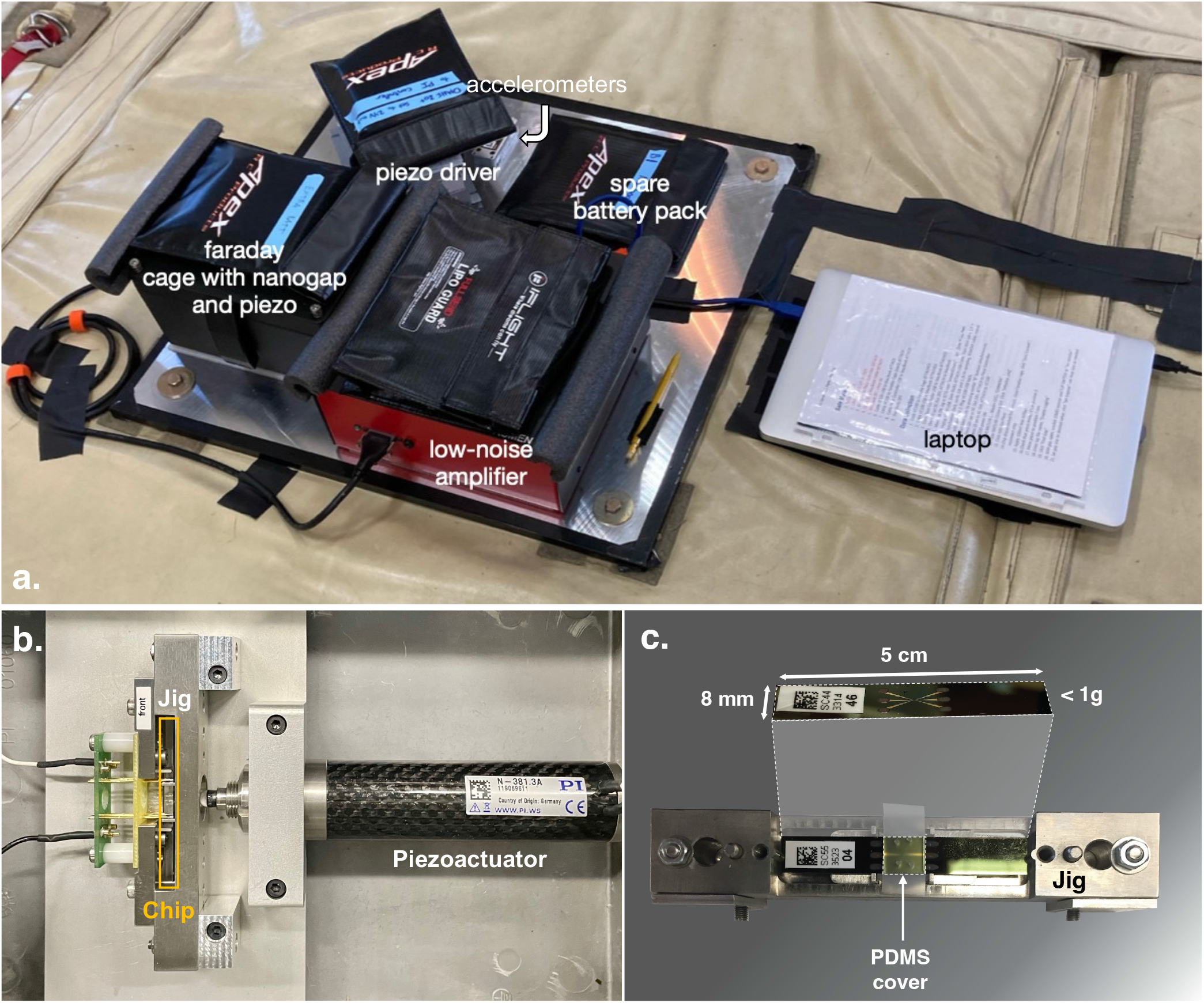
Next-generation ELIE instrument prototype. (a) ELIE system consisting of a laptop controlling the low-noise amplifier that, in turn, supplies and controls voltage to and within a Faraday cage that encloses (b) a piezoactuator, and a jig structure holding the (c) nanogap chip.

### ELIE Instrument Operations and Testing

The chip, positioned within a Faraday cage, was mounted on a three-point bending mechanism using a jig. It was rinsed with 10% ethanol as a wetting agent and mechanically bent with the integrated piezoactuator to create an atomically sharp gap. A 10 kΩ resistor was connected in series to prevent over-current breakdown during connection. After forming the gap, the resistor was disconnected to allow the current measurement to reflect gap electron tunneling conductance. Gap characterization was performed in a dry state before the sample measurement. L-proline, purchased from Sigma (81709), was then used to prepare a 10 *μ*M solution in nuclease-free water with 1 mM phosphate buffer (Sigma P3619) at pH 7.4 and 25 °C. A 0.1 mL volume was introduced through the microchannel in the PDMS, and current was recorded during the 1-hour, 22-minutes, and 44-seconds flight at an applied voltage of 100 mV with a 4 MHz sampling rate. Ground testing, lasting 1-hour, 7-minutes, and 30 seconds, utilized a solution with the same L-proline concentration and ELIE system parameters. Adjustments to the gap were achieved by bending the chip with the piezoactuator, resulting in changes relative to the vertical motion. Current, monitored by VC100, accounted for offset error, ionic current, and tunneling current. During analysis, the baseline was zeroed to emphasize current differences when a molecule occupied the gap.

### Signal Processing and Event Detection

Recorded data was pre-processed by applying a low-pass filter at 10 kHz and down-sampling to 100 kS/s to mitigate electrical noise and reduce data storage requirements, respectively. Subsequently, baseline adjustment was performed by computing the median of the current data. Event detection was accomplished using Trans-analyzer, an open-source MATLAB GUI-based package for nanopore signal analysis developed by Calin Plesa at Deft University of Technology [26]. We used a 1000-point moving average window for baseline detection to ensure an accurate representation of the local baseline in the presence of fluctuating values. The analysis employs a thresholding algorithm for event detection, where events are identified if they exceed a specified threshold above the local baseline level. The determination of this threshold involves multiplying a peak detection factor by the root mean square (RMS) noise level. A peak detection value of 6 was chosen to minimize noise spikes while maximizing L-proline events capture. To determine the noise level, the moving standard deviation histogram peak method was used. This method calculates standard deviations within 1000-point windows and creates a histogram of these values to pinpoint the standard deviation of the baseline (noise), estimated as a peak in the histogram. To improve event detection accuracy and stabilize baselines in our 1000-point moving average window, we applied an iterative detection method three times. In each iteration, the algorithm generated a new trace, replacing the detected event’s duration with the local baseline value at the event’s start, effectively decoupling the moving average calculation from the influence of prior events.

### Acceleration data measurements, processing, and flight profile segmentation

The two accelerometers (metal-body Slam Stick X; Mide Technology Corp.) were mounted next to ELIE on a common baseplate, using double-sided tape (3M 950), to accurately measure vibrations across a wide range of frequencies with minimal distortion. The flight profile was segmented as described in Carr *et al*. 2018 from the acceleration data collected [27]. In short, all the flight phases (defined by the different *g* levels) were first identified using a change point detection, as implemented by the MATLAB *FindChangePts()* function. Then, to categorize flight periods by stable gravitational levels and detect transitions, a secondary change point detection method was employed, involving linear slope analysis within 10-second intervals near each change point. This process effectively separated the flight data into regions of rapid “transition” and more stable periods. These stable, “non-transition” flight periods were further classified into distinct phases of “parabola,” “hypergravity,” and “other”. The classification process involved first categorizing all periods longer than 100 seconds as “other”, which represented stabilized level-flight periods (gravitational forces between 0.9*g* and 1.3*g*). The remaining periods were subsequently segmented based on gravitational levels, with “parabola” assigned to periods featuring gravitational forces less than or equal to 0.9*g*, and “hypergravity” to forces exceeding 1.3*g*.

### ELIE and Acceleration Data Integration

Each event was assigned to one of the flight phases (parabola, transition, hypergravity, other) based on the periods.*txt* file produced by prior analysis. Following this, outliers in the data were detected using the boxplot method, which divides the data into lower and upper quartiles, encapsulating the middle 50% of the data within the interquartile range (IQR). Outliers are identified as data points that fall 1.5 times or more above the upper quartile or below the lower quartile (Supplemental Figures 1 and 2). Comparisons between flight phases were subsequently conducted using features exclusively from true events, including tunneling current (average current level within a single event) and dwell time (duration) for each event. Furthermore, the gap size during each of these events was estimated by employing the theoretical relationship of the baseline current (*I*) and the gap distance (*d*). The gap distance is determined by the exponent<u>ial fu</u>nction of *I* ∼ exp(*βd*) with the decay constant 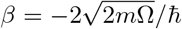, where *m*, Ω, and 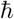 are the electron mass, the work function of gold (Au), and the reduced Planck’s constant, respectively [25].

## 3. Results

### Parabolic Flight and G-levels Achieved

Flight operations were executed aboard G-Force One®, Zero Gravity Corporation, a Boeing 727-200F aircraft designed to conduct parabolic maneuvers. A series of 20 parabolic maneuvers were conducted, comprising four distinct sets, each consisting of five individual parabolas (see Figure 3a). In the first set, the targeted accelerations, in sequence, were Mars *g*, Mars *g*, Lunar *g*, Lunar *g*, and zero *g*; the parabolas within this set are clearly discernible in Figure 3b at 1000-1500 s. All subsequent parabolas were exclusively aimed at achieving zero *g* conditions. The flight profile was segmented into discrete phases, including “transition,” “parabola,” “hypergravity,” and “other”, based on accelerometer measurements as described by Carr *et al*. 2018 (Figure 3c) [27]. The mean duration of the flight parabolas were 18.12 ± 1.18 s (zero *g*, N = 17), 22.02 ± 1.42 s (Lunar *g*; N = 2), and 29.27 ± 1.23 s (Mars *g*; N = 2), while the corresponding achieved gravitational forces were 0.041 ± 0.006 *g* (zero *g*), 0.154 ± 0.014 *g* (Lunar *g*), and 0.370 ± 0.033 *g* (Mars *g*). Each flight phase was validated through data analysis obtained from the two accelerometers positioned on the flight baseplate, demonstrating complete alignment in the acceleration data (Supplemental Figure 3). Vibration data was not collected on the ground; vibration measurements from a prior flight suggested about a factor of 6 between ground and flight [28].

**Figure 3.**
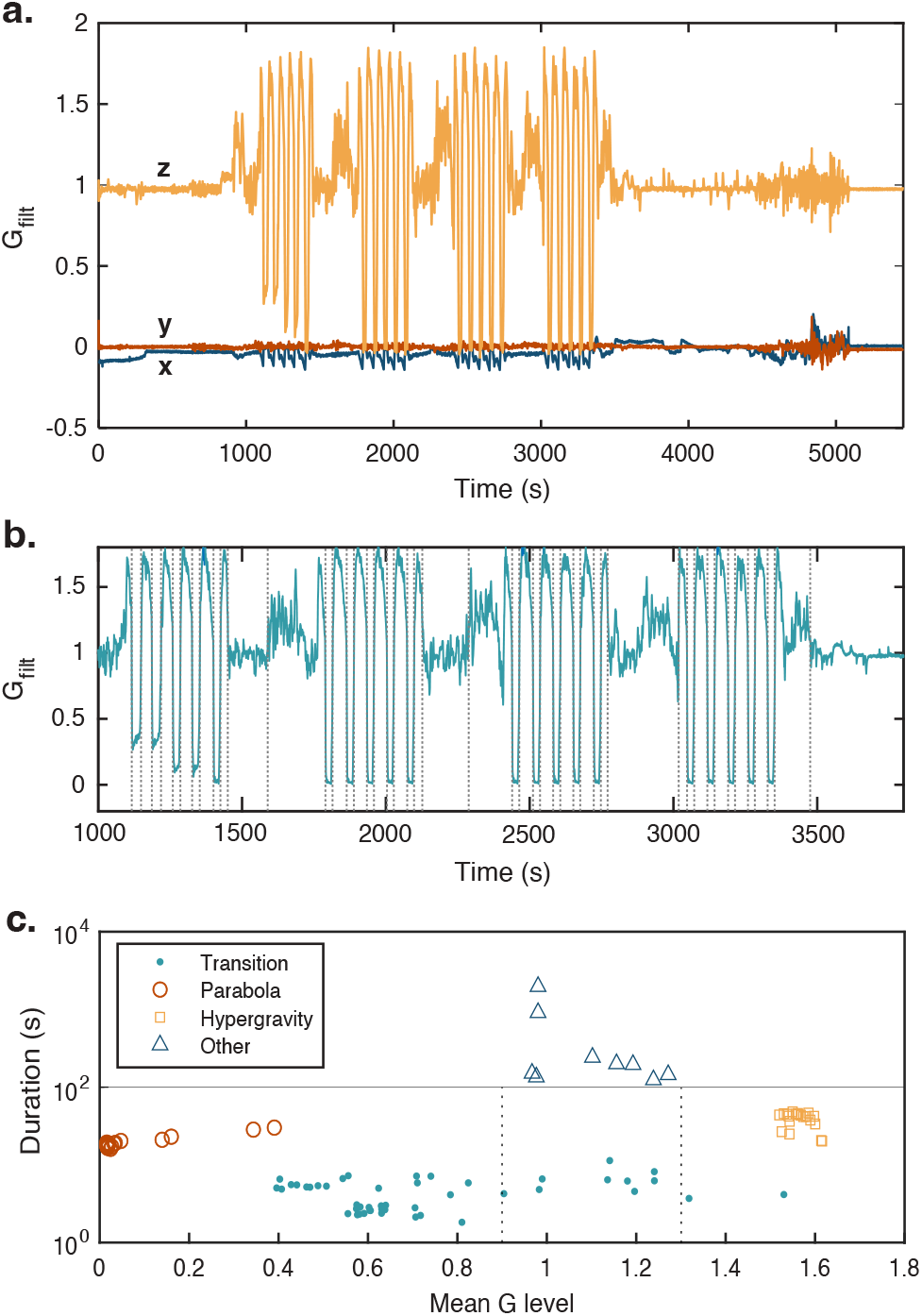
Analysis of parabolic flight acceleration data: filtering, phase segmentation, and classification. (a) Measured accelerations after filtering. (b) Change points, delineated by the vertical lines, for mean *g* levels. (c) Categorization of transition and non-transition periods by duration and mean *g* level.

### Detecting Amino Acid L-proline across various G-levels

Single-molecule detection of L-proline was demonstrated with the next-generation ELIE system using a 10 *μ*M L-proline aqueous solution and an automatically adjusted gap size between the electrodes. ELIE data were collected for a total duration of 82 minutes during flight and 67.5 minutes on the ground. Given the inherent noise in the data, only 5.2 minutes of flight data and 30.2 seconds of ground data were of sufficient quality for subsequent event detection analysis. The flight data encompassed all flight phases experienced during the parabolic flight, including two zero *g* parabolas, one Martian parabola, one Lunar parabola, ten hypergravity periods, seven transition periods, and four level-flight periods. The parabola phase had a mean period duration of 22.09 s ± 4.32 s, while the remaining phases exhibited variable durations, with an average duration of 36.02 s ± 9.20 s for the periods in the hypergravity phase, an average duration of 4.60 s ± 1.88 s for the periods in the transition phase, and an average duration of 835.62 s ± 879.56 s for the periods in the other phase. For the event-level analysis, the events detected during the transition phase were placed under the “other” category due to their varying gravitational levels. Examples of current–time profiles obtained from these time periods during flight and ground are shown in Figure 4.

**Figure 4.**
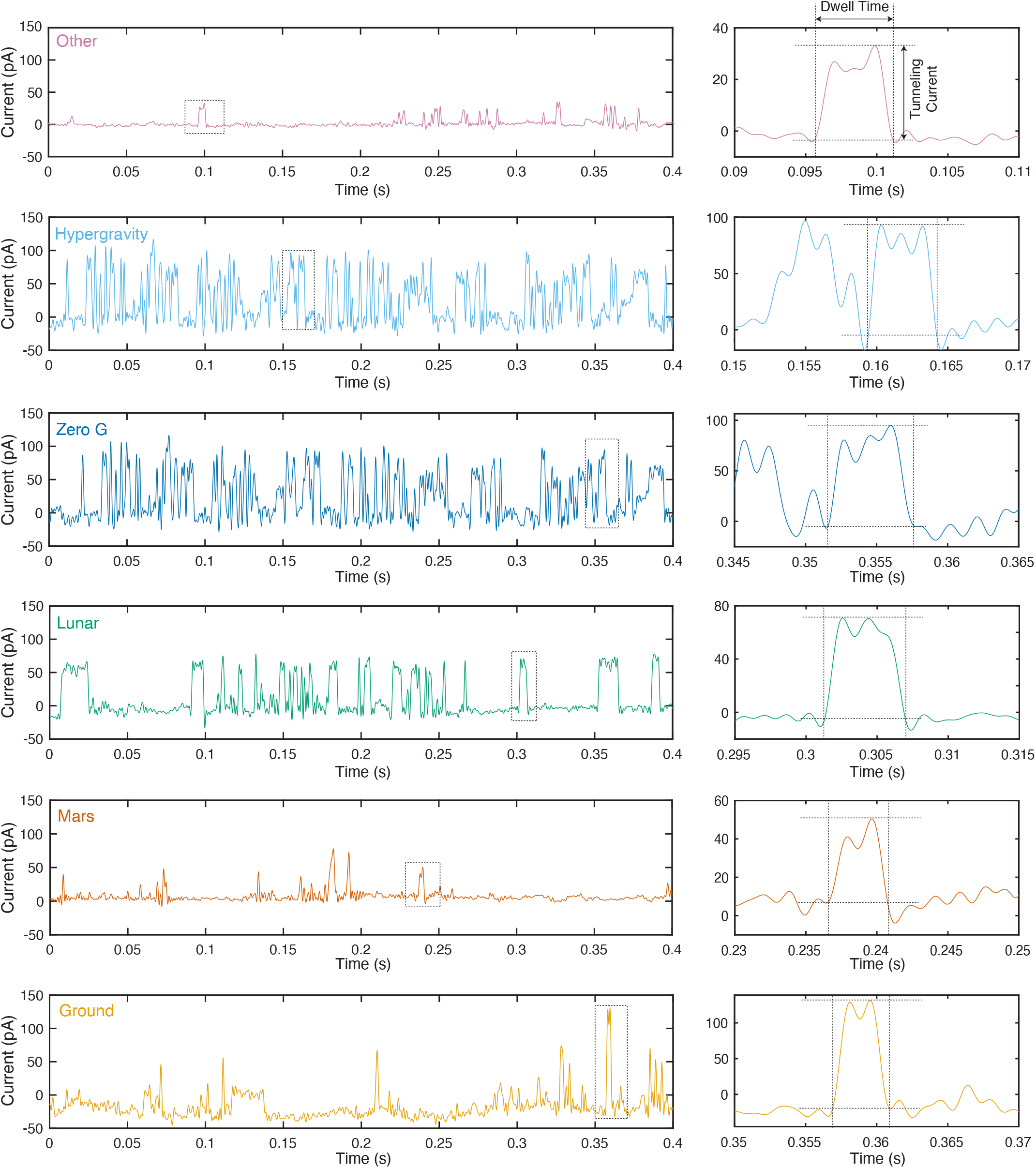
Current-time profiles of L-proline events across different flight periods and during a period of the ground experiment. The dash-lined boxes in the left plots correspond to the zoomed-in plots on the right side of the figure. Events are characterized by an average tunneling current and the duration of the current (dwell time).

Following outlier identification, the number of events detected per parabola was as follows: 156 for the first zero *g* parabola, 663 for the second zero *g* parabola, 332 for the Martian parabola, and 812 for the Lunar parabola. On average, 491 ± 300 L-proline events were detected per parabola period, 460 ± 343 per hypergravity period, and 435 ± 564 per the other transition and level-flight periods. The data collection yielded event rates of 21 ± 1 events per second for zero *g* parabolas, 12 events per second for the Martian parabola, 39 events per second for the Lunar parabola, 13 ± 1 events per second for the hypergravity phase, and 1 ± 1 event per second for the “other” phase. Notably, the subcategories within the parabola phase happened to have the highest event rates but also the shortest in total duration during the parabolic flight, accounting for just 1.07%, 0.81%, and 5.32% of the overall parabolic flight duration for Mars parabolas, Lunar parabolas, and zero *g* parabolas, respectively. This may simply reflect ongoing experiment monitoring and adjustments during the non-parabola portions of flight.

### Event Level Analysis

The tunneling current and dwell time histograms depicted in Figure 5a and 5b provide insight into the distributions of the detected events. Figure 6a provides a comprehensive overview of event statistics, including the number of events, average tunneling current, and average dwell time per parabolic flight phase. This table extends our analysis by incorporating data from the parabolic flight into the context of prior ground experiments involving L-proline detection using the AXN system and the first-generation ELIE prototype [21, 24]. The distribution of average conductance versus average dwell time for each parabolic flight phase, along with the results from the aforementioned ground experiments, is visually presented in Figure 6b. This figure suggests that the gap size produced by the piezo actuator was close to 0.70 nm during the parabolic flight when compared to the AXN data. Note that the estimation of the gap size, based on the current signal of the baseline, suggests average gap size per flight phase ranging from 0.33 nm to 0.44 nm during periods of good quality data acquisition. However, this analysis is under the assumption of single gold atom-to-atom contact, as illustrated in Figure 1. These findings deviate from the established relationship between gap distance and tunneling current previously outlined by in Tsutsui *et al*. 2008, where a larger gap distance corresponds to lower tunneling currents and, consequently, lower conductance [25]. This may be due to uncharacterized changes in gap geometry, as summarized in the discussion.

**Figure 5.**
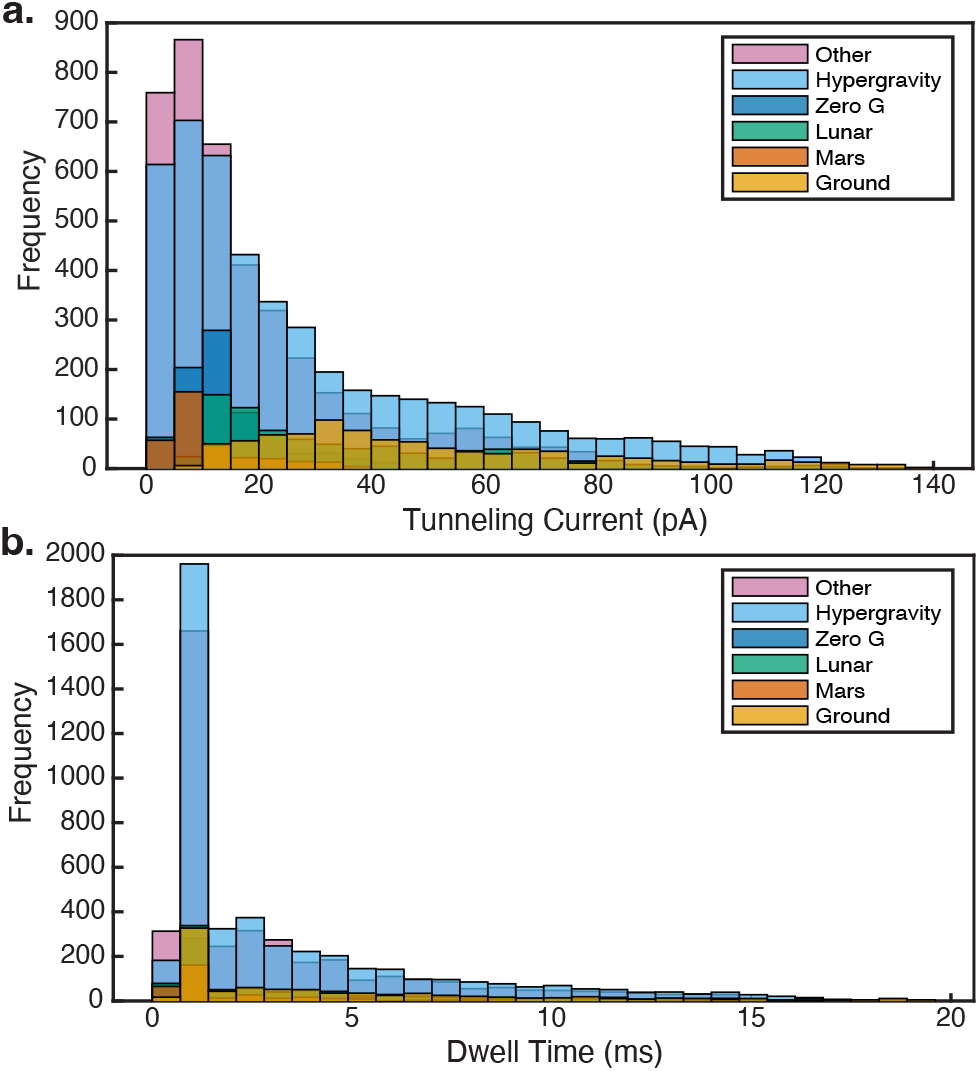
L-proline Event Level Analysis across Flight Phases and Ground. (a) Average tunneling current and (b) dwell time histograms of the L-proline events detected during each flight phase and ground analysis.

**Figure 6.**
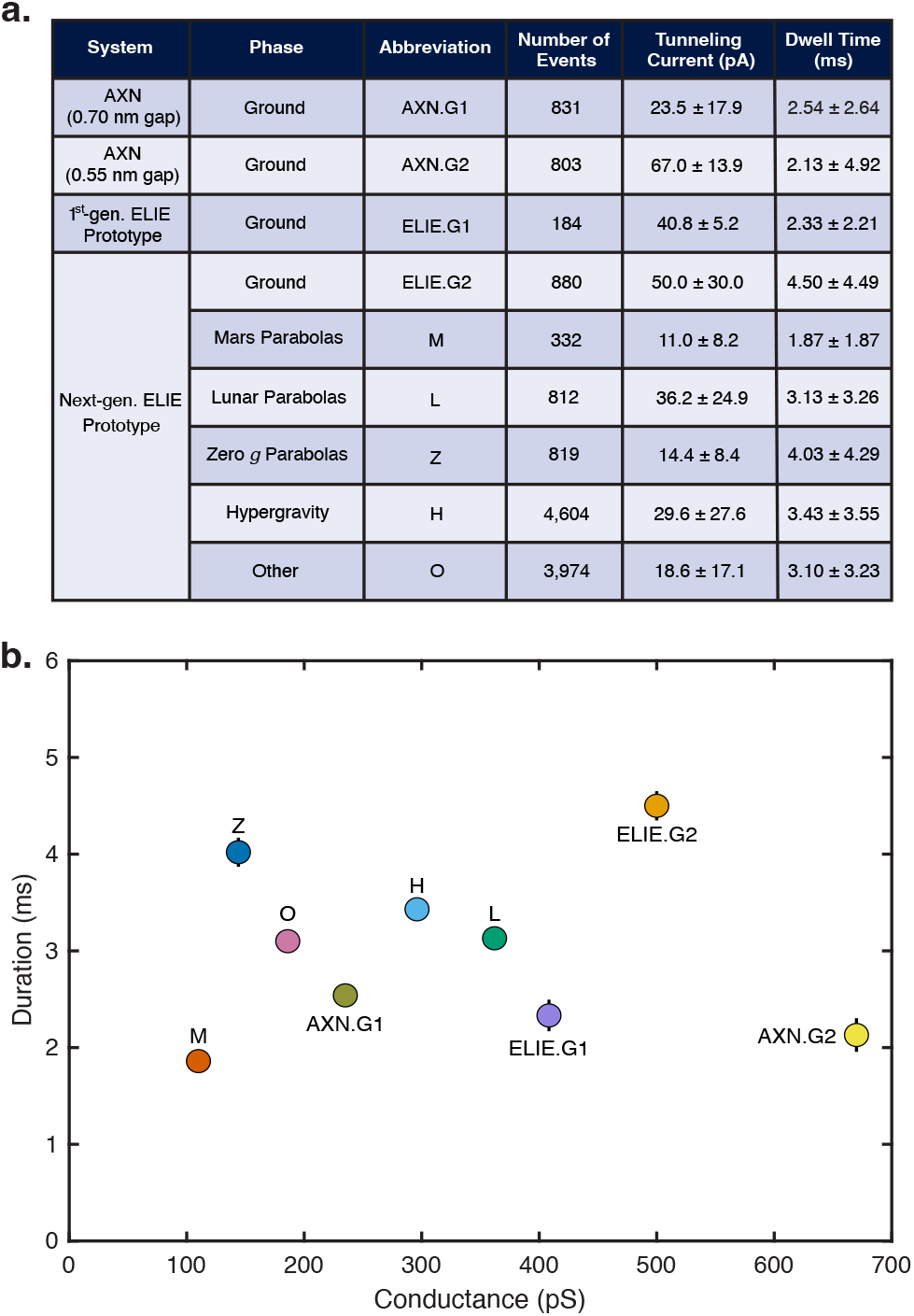
L-proline Event Level Analysis across Flight Phases and Ground. (a) Table summary of single-molecule average tunneling current and dwell time for L-proline across the different flight phases experienced by ELIE during the parabolic flight, and ground experiments performed by the next-generation ELIE prototype, the first-generation ELIE prototype [24], and the AXN system [21]. (b) Plot of single-molecule mean conductance and mean dwell time of L-proline events obtained during the parabolic flight phases and the ground experiments. Refer to the table for the designation of each phase.

## 4. Discussion

### Limitations and Future Developments

The next-generation ELIE prototype’s ability to produce reliable event detection is hindered by the high levels of noise experienced during operation. With the average tunneling current measurements being on the scale of tens of picoamperes, significant noise in that regime can lead to issues such as false detections or events being mistakenly filtered out as noise. To address this problem, noise reduction within the ELIE prototype is essential. One potential source of noise is the power supplies used to run the instrument during flight (Omni20 battery pack), which provided either DC voltages up to 24V or AC power that utilized a switching regulator. Additionally, we empirically observed a change in noise when the low-noise amplifier cable was moved, indicating external coupling of noise into the system.

The noise may have also been influenced by surrounding electronics and environmental conditions, such as temperature, pressure, and the vibrational environment. Measurements taken on the ground for this experiment exhibit more noise than measurements taken during flight, which seems contrary to what would be expected given the vibrational environment and gap decaying that was experienced during flight. This could be due to certain external noise sources being more unstable or unpredictable than others, which can be better characterized. It could also be attributed to the use of a wall power supply for ground measurements while battery power was utilized for flight measurements.

Another limitation relates to nanogap stability and the limitations of conducting the parabolic flight. The nanogap chip in the instrument could not be changed before or during the flight, and the gap had to be pre-formed on the ground the prior day. Therefore, the chip had been sitting in fluid overnight before measurement. Over time, it is likely that the gap decayed, changing from a sharp single gold-gold atomic contact to a more-blunt multiple gold atom contact. This likely led to an underestimate of gap size, and may also have resulted in an unstable gap during the system’s startup in flight, as evidenced by the lower number and duration of detected events during the Mars parabola, the first flight parabola achieved, which distinctly stands out as an outlier compared to the events detected during the other flight phases (Figure 6a, Supplemental Figure 2). Evidence for this also included changes in the relationship between piezo position and estimated gap size (data not shown). These presumed geometry changes may also explain the observed variations in the conductance and duration between phases of flight (Figure 6b).

Several measures are underway to reduce the impact of noise and improve nanogap characterization. Using a cleaner power supply to energize the prototype is one such measure. Using the same power source for both flight and ground experiments will also be important to improve the interpretation of the data. Reducing or eliminating external coupling of noise, for example, cable shielding or complete Faraday cage protection, is also important. Future work could also probe the impacts of vibrational and temperature changes and their association with noise. Additionally, it is important to characterize the geometry of the nanogap. This can be done by monitoring the current during gap formation to determine changes in resistance during formation over time, indicative of gap size and shape. With a better knowledge of gap size, the estimation of gap distance can be more accurately characterized.

In addition to high noise levels, the next-generation ELIE prototype had areas of dead space in the Faraday cage and a mass over 8 kg which could prove prohibitive to a spaceflight instrument. To remedy this, new parts can be chosen and the Faraday cage redesigned in future ELIE prototypes that reduce these metrics. Additionally, the next-generation ELIE prototype currently uses a manual sample loading system. Though this is effective for bench testing, this greatly limits the volume of sample that can be analyzed by the instrument and would limit the instrument to only be used in situations where human interaction is possible. Therefore, an automatic sample delivery system must be implemented into future ELIE prototypes to allow for a larger volume of sample to be able to be analyzed by a singular nanogap chip. With an automated sample delivery system and the automation in the next-generation ELIE prototype, a fully automated prototype can be developed.

### Applications

The ELIE prototype is currently at a TRL 2. However, with further development, ELIE aims to become a fully automated system that can take in a preconcentrated sample and detect and identify any single molecules present in the sample accurately. This functionality would allow ELIE to discern an array of molecular entities, ranging from amino acids to informational polymers. Such a capability holds immense significance for the exploration of celestial bodies like Europa and Enceladus, where the nature of potential biosignatures remains enigmatic, as does the form life may take if it exists. Consequently, ELIE stands as a compelling candidate for integration into upcoming missions such as the Orbilander mission to Enceladus, scheduled for launch in 2038. Its potential deployment on the lander would facilitate real-time, *in-situ* identification of potential biosignatures [29].

## 5. Conclusions

Ongoing efforts are aiming to enhance the feasibility of the Electronic Life-detection Instrument for Enceladus/Europa for *in-situ* life-detection missions to Ocean Worlds. ELIE’s ability to detect single molecules, such as the amino acid L-proline, under different gravitational conditions, including microgravity and reduced gravity (e.g., Mars, Lunar, and Europa), marks a significant step forward in our quest to make solid-state nanogaps viable for molecular biosignature detection. The integration of an automated piezo actuator has enabled real-time gap control, addressing previous limitations. However, many challenges remain, including power supply noise and chip-related noise, which impact the quality of data collected. Future developments aim to reduce size and mass, lower noise levels, and further automate the system to reach TRL 4. With continued advancements, ELIE has the potential to become an autonomous and highly sensitive single-molecule detection instrument for deployment across our solar system, aiding in the search for life beyond Earth.

## Acknowledgments

We thank the MIT Media Lab Space Exploration Initiative for providing the parabolic flight, which was supported by NASA Award 80NSSC21M0012 to M.T.Z. This work was supported by NASA awards 80NSSC19K1028 and 80NSSC22K0188 to C.E.C.

## Appendix

### 1. Supplemental Figures

**Supplemental Figure 1:**
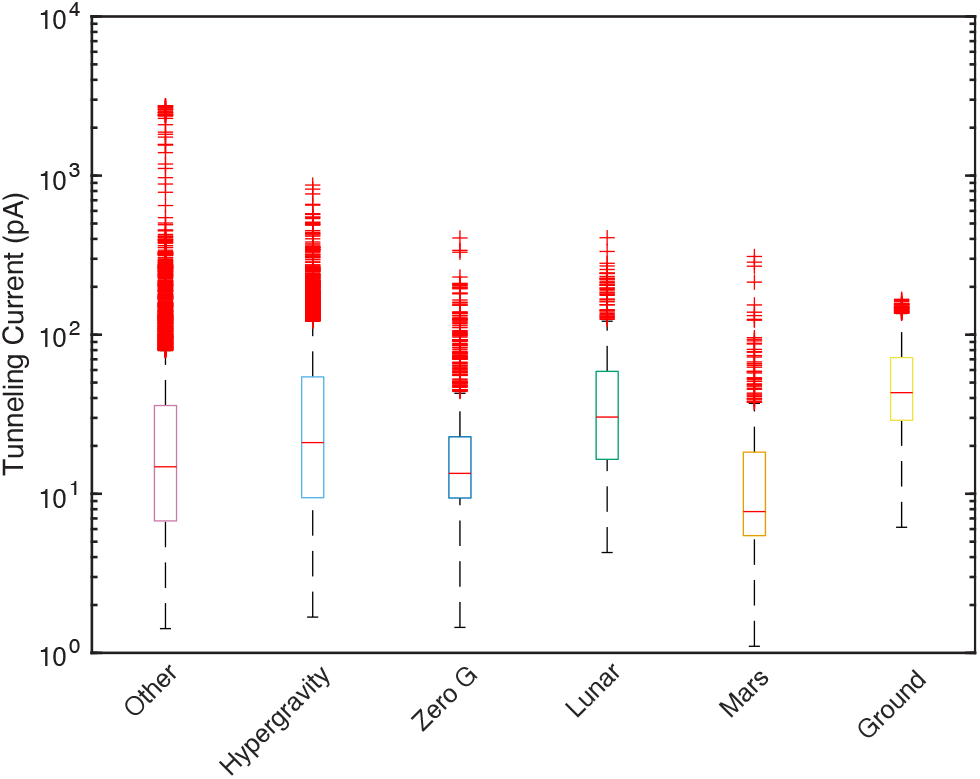
Outlier identification using the IQR method based on tunneling current of detected events.

**Supplemental Figure 2:**
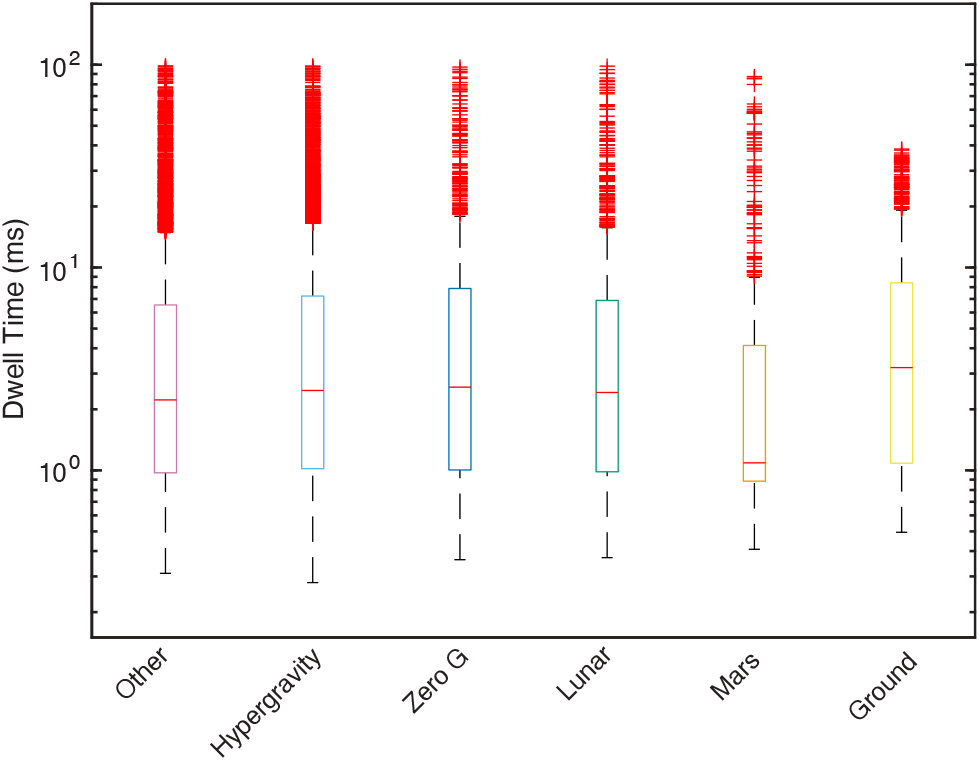
Outlier identification using the IQR method based on dwell time of detected events.

**Supplemental Figure 3:**
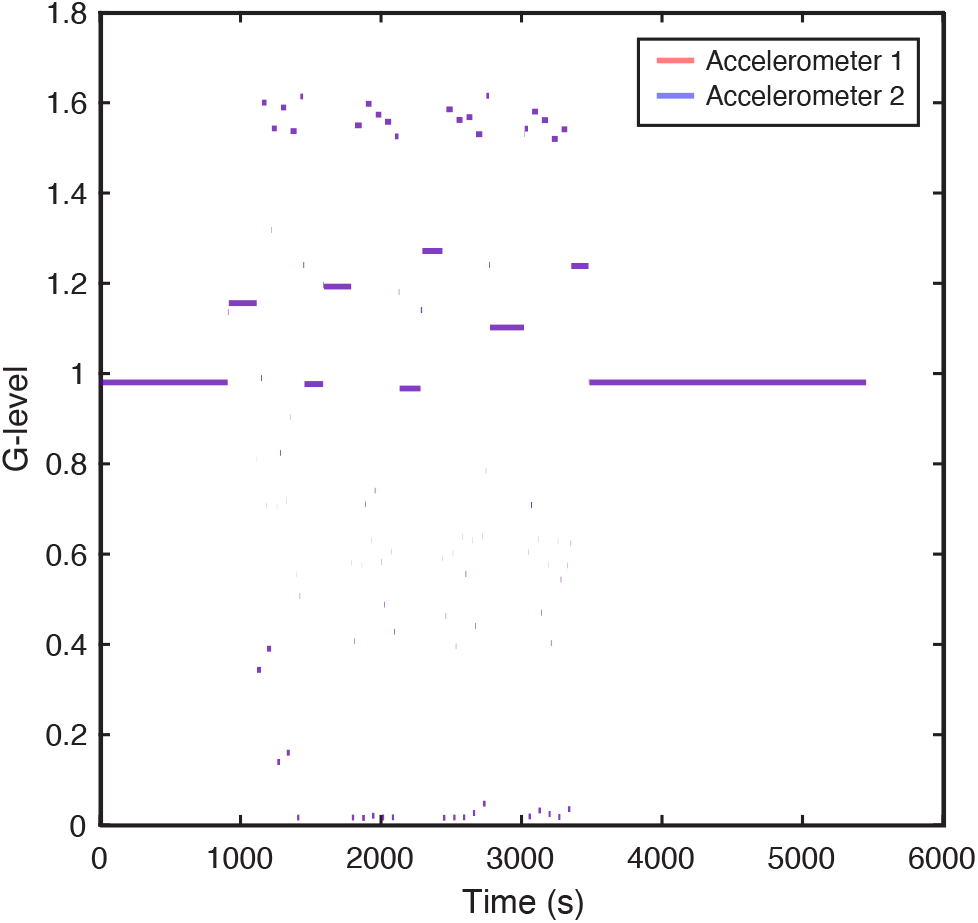
Timeline of flight phases recorded by onboard accelerometers.

## Notes

### Competing Interest Statement

The authors have declared no competing interest.

